# Reinforcement influences the ability of cryptic female choice to exert conspecific sperm precedence in hybridizing Atlantic salmon (*Salmo salar*) and brown trout (*Salmo trutta*)

**DOI:** 10.64898/2026.05.08.723816

**Authors:** Connor P. Hanley, Ranjan Wagle, Sarah J. Lehnert, Craig F. Purchase

## Abstract

Conspecific sperm precedence via cryptic female choice is a post-ejaculatory selection process that reduces hybridization, and can be pronounced in sympatric species. In their native Europe, Atlantic salmon (*Salmo salar*) and brown trout (*Salmo trutta*) exert conspecific sperm precedence under heterospecific sperm competition, which is at least partially enabled by female reproductive fluid. We examined post-ejaculatory selection of both species in Newfoundland, Canada, where Atlantic salmon evolved in absence of brown trout, but now experience hybridization threats due to anthropogenic introductions. Using split-ejaculate and split-clutch in-vitro fertilizations we evaluated whether allopatric evolution has relaxed this selection in Atlantic salmon, and found that they had no ability to bias paternity towards conspecific males, whereas naturalized brown trout retained a strong ability to do so. Female reproductive fluid influenced this, as when fluid associated with a species’ eggs was swapped, hybridization increased. In the artificial situation of no female reproductive fluid during sperm competition, paternity changed dramatically, but sperm swimming performance did not predict it. Our findings contribute to understanding the evolution of cryptic female choice and how the mechanisms of reproductive isolation can be reinforced through sympatry, while also highlighting a new potential conservation concern for North American Atlantic salmon.

## Introduction

Sexual selection “arises from fitness differences associated with nonrandom success in the competition for access to gametes for fertilization” (Shuker and Kvarnemo 2021). In the pre-ejaculatory diploid phase of sexual selection (Parker, 2014; Purchase et al., 2021), mate choice allows individuals to pick preferred mates that can be advantageous for their own offspring (Darwin, 1871). Females invest more energy than males into each individual gamete (Clutton-Brock, 1991; Schärer et al., 2012), consequently limiting the number of gametes and mating opportunities, and as a result females are typically more selective than males in choosing mates. It is essential that they identify and avoid mating with incompatible males, such as those of different species, and instead select the best males of conspecific species, often based on secondary sexual characteristics such as size, color, smell, sound, and ornamentation (Edward, 2015; Ferkin, 2018; Hanson & Smith, 1967; Milinski & Bakker, 1990; Searcy, 1992). Males produce numerous tiny sperm at a relatively low cost (Hayward & Gillooly, 2011; Parker, 1997) and allocate most of their reproductive energy towards creating mating opportunities through competition with other males (Candolin, 1999; Petersson et al., 1999) and courting females (Jamieson, 2003). Post-ejaculation, sexual selection continues in the haploid phase as sperm of different males compete to achieve fertilizations of a female’s eggs (Parker, 2020). The outcome of sperm competition is mediated by sperm quantity and quality (Crean & Marshall, 2008b; Gage et al., 2004; Gage & Morrow, 2003; Kekäläinen et al., 2015) and cryptic female choice (Birkhead, 1998; Eberhard, 1996; Firman et al., 2017), which biases fertilization in sperm competition towards a certain male. Cryptic female choice can reinforce mate choice from the earlier pre-ejaculatory stage to the same potential father (see Figure 1a: ‘within-generation haploid-diploid reinforcement’), including biasing paternity towards conspecific males when sperm from heterospecific individuals are present. This process is referred to as conspecific sperm precedence (Howard et al., 1998) and can be an important step in reducing hybridization (Purchase et al. 2024).

**Figure 1:**
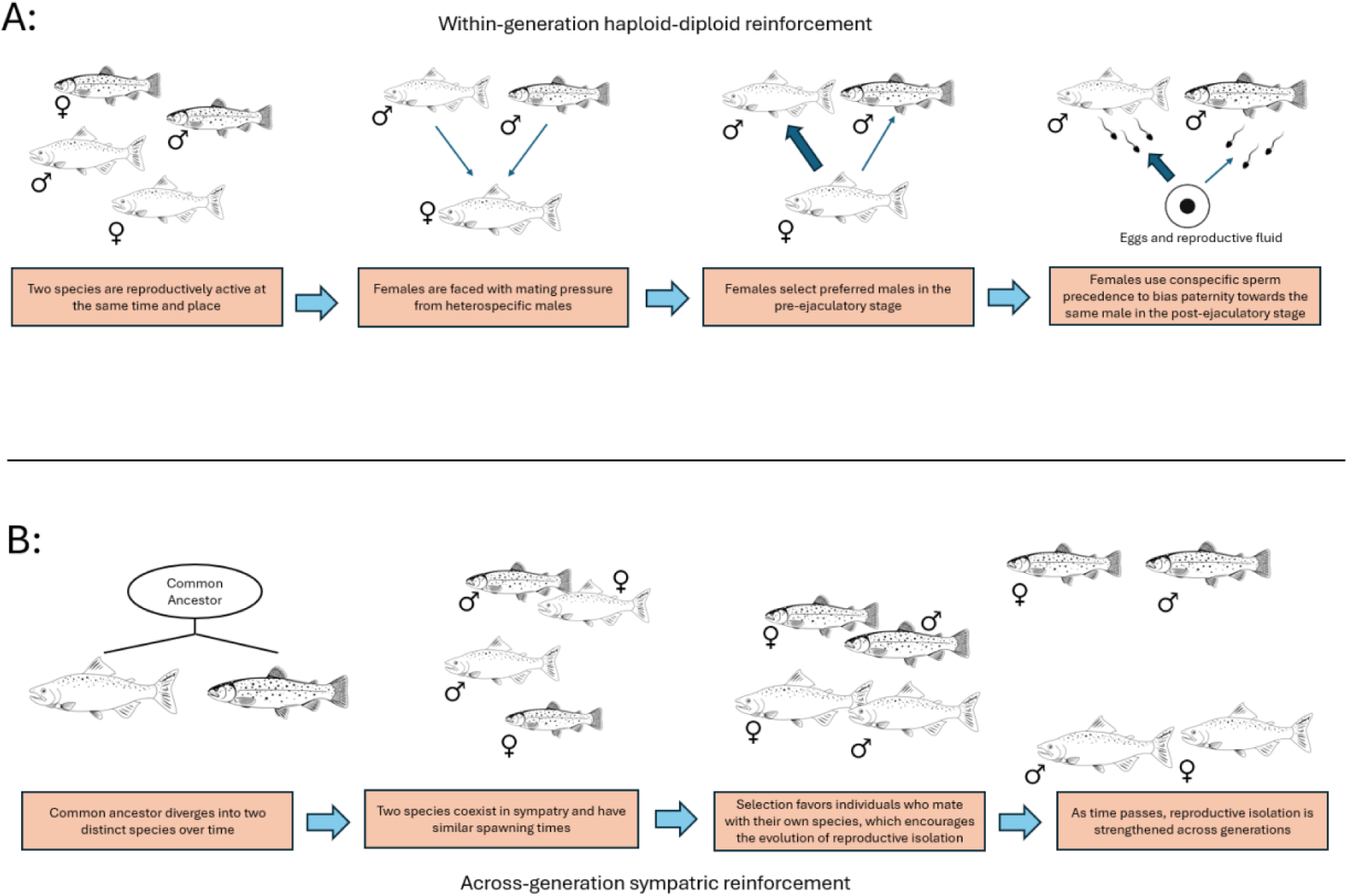
Defines ‘reinforcement’ in two contexts related to reproduction. The key question addressed in this study is whether the strength of B influences the strength of A. As far as we are aware, no previous study has used both of these concepts in tandem. (A) Sexual selection “arises from fitness differences associated with nonrandom success in the competition for access to gametes for fertilization” (Shuker and Kvarnemo 2021). If pre-ejaculatory and post-ejaculatory selection align to select the same father ‘within-generation haploid-diploid reinforcement’ occurs between the haploid and diploid phases of the biphasic life cycle and strengthens mate choice among the same individuals. Examples: (House et al., 2016; Howard et al., 1998; Paczolt & Jones, 2010; Parker, 2009; Rebar et al., 2011, 2019) (B) ‘Across-generation sympatric reinforcement’ strengthens reproductive isolation between related species through time via a positive feedback loop. Examples: (Arce-Valdés et al., 2024; Hopkins, 2013; Moran et al., 2018; Noor, 1999; Nosil et al., 2003; Pfennig & Rice, 2014; Rundle & Schluter, 1998; Servedio, 2004; Servedio & Noor, 2003; Stre et al., 1997)

Hybridization occurs when individuals from two genetically distinct populations successfully breed with one another and produce offspring (Abbott et al., 2013; Harrison & Larson, 2014). Hybridization can have varying effects on offspring performance (Adavoudi & Pilot, 2022; Burke & Arnold, 2001), potentially increasing its fitness through hybrid vigor (Hubbs, 1955). However, when animals hybridize the resulting offspring are usually less fit, which presents a net negative impact on the parents’ fitness. Outbreeding depression lowers fitness (Barton, 2001; Birchler et al., 2006) through either increased mortality (Puigcerver et al., 2014; Saulsberry et al., 2017) or decreased reproductive success (Bhattacharyya et al., 2013; Sweigart et al., 2006). A hybrid individual that fails to reproduce is not only a waste of the energy that the parents put into creating it, but also a source of competition for purebred half-sibling offspring. Due to investing more into each individual gamete and having limited mating opportunities, the consequences of each hybrid mating are far larger for females compared to males (Purchase et al. 2024). Therefore, females have a greater pressure to avoid heterospecific fertilizations through mechanisms that reduce heterospecific matings (pre-ejaculatory) and fertilizations (post-ejaculatory). Over generations, when closely related species exist in sympatry, these mechanisms can evolve and reinforce species boundaries (Dobzhansky, 1951; Hopkins, 2013; Howard, 1993; Servedio, 2004; Servedio & Noor, 2003), while under allopatric conditions such reinforcement does not occur, and these mechanisms might be less effective (see Figure 1b: ‘across-generation sympatric reinforcement’).

Hybridization is more common when historically allopatric populations become contemporarily sympatric due to anthropogenic introductions, and invasive species often have the potential to interbreed with genetically distinct native species (Allendorf & Leary, 1988; Kovach et al., 2015). There is evidence to indicate that when ‘across-generation sympatric reinforcement’ (Figure 1b) is absent, hybridization increases on secondary contact due to weaker pre-ejaculatory mate choice. For example, allopatric females can show less mate discrimination against heterospecific males in toads (Pfennig & Rice, 2014), birds (Stre et al., 1997), and fish (Rundle & Schluter, 1998) compared to sympatric populations. In fish, allopatric males compete against heterospecific males at higher frequencies and select heterospecific females more often than sympatric males do (Gabor & Ryam, 2001; Moran et al., 2018; Moran & Fuller, 2018). Less information is available on how ‘across-generation sympatric reinforcement’ (Figure 1b) influences the strength of post-ejaculatory sexual selection. Evidence has shown that sympatric populations of flies have a greater ability to exert conspecific sperm precedence than their allopatric counterparts (Castillo & Moyle, 2019). In the same study however, compared to allopatric females, sympatric females in other populations had a weaker ability to use intra-population cryptic female choice, indicating that investment into preventing males of different species from achieving fertilization may in turn reduce the opportunity for post-ejaculatory sexual selection within a population. There is a dearth of information on whether the degree of ‘across-generation sympatric reinforcement’ (Figure 1b) – i.e. positive feedback through generations from being sympatric, influences the strength of ‘within-generation haploid-diploid reinforcement’ (Figure 1a) via cryptic female choice. Such a pattern might be predicted if there were costs to maintain reproductive isolation via cryptic female choice when it was no longer needed. If a previously sympatric species was separated from the closely related heterospecific counterpart, would the strength of conspecific sperm precedence via cryptic female choice degrade?

Salmonid fishes are an excellent group to address this question (Purchase, 2022). Cryptic female choice is easily manipulated as it is mediated outside the body by female reproductive fluid (hereafter = ovarian fluid), which is stored and released alongside eggs (Rosengrave et al., 2008; Urbach et al., 2005; Yeates et al., 2013). Salmonids are polygamous (De Gaudemar, 1998), readily able to hybridize (Chevassus, 1979), have males that use alternative reproductive tactics (Fleming, 1996), and have varying levels of sympatric reinforcement (Lantiegne & Purchase, 2023; Purchase, 2022; Yeates et al., 2013). Atlantic salmon (*Salmo salar*) and brown trout (*Salmo trutta*) are two relatively well-studied sister species that are morphologically similar but are genetically distinct (Lecaudey et al., 2018; Lubieniecki et al., 2010; Pendas et al., 1995). In Europe, both species are native and are often reproductively active in the same places and can spawn at the same times, and thus have co-evolved in sympatry. Both species demonstrate complete gamete compatibility, however hybrid offspring are always infertile by the second generation (Chevassus, 1979; Purchase et al., 2024) and because of this there should be strong selective pressure to avoid heterospecific fertilizations (Purchase et al., 2024). There is little information regarding pre-ejaculatory mate choice mechanisms to avoid hybridization, but a study has been done on post-ejaculatory conspecific sperm precedence. Research in Europe demonstrates the complete interfertility of these species when fertilizing without sperm competition, but when sperm of the two species compete, males that are conspecific to the female achieve a much higher portion of paternity compared to heterospecific males (Yeates et al., 2013). This is at least partially enabled by the ovarian fluid, as when the ovarian fluid associated with a female’s eggs is switched with the other species’ ovarian fluid, heterospecific males achieve a higher proportion of paternity (Yeates et al., 2013).

European populations of Atlantic salmon and brown trout that have evolved in the presence of ‘across-generation sympatric reinforcement’ (Figure 1b) show a strong ability to exert conspecific sperm precedence (Yeates et al., 2013), complementing pre-ejaculatory mate choice with ‘within-generation haploid-diploid reinforcement’ (Figure 1a) after ejaculation.

North American Atlantic salmon have been separated from their European counterparts for over 600,000 years (Lehnert et al., 2020) and thus likely have been separated from brown trout for at least that long. During this time North American Atlantic salmon would not have co-evolved with brown trout and thus do not have historical ‘across-generation sympatric reinforcement’ (Figure 1b), which might have led to a loss of ‘within-generation haploid-diploid reinforcement’ (Figure 1a). Brown trout are an invasive species in North America, being first established in Newfoundland in 1883 (Hustins, 2007). Since their introduction, brown trout have invaded watersheds that contain Atlantic salmon (MacDonald et al., 2024; Westley & Fleming, 2011).

However, the spread of brown trout in Newfoundland is unusually slow when compared to invasions in other places around the world (MacDonald et al., 2024; Westley & Fleming, 2011), and while this could be due to a variety of reasons, the fertilization of brown trout eggs by Atlantic salmon sperm could be a significant factor in preventing trout from establishing sustainable populations (Lennox et al., 2025; MacDonald et al., 2024; Purchase et al., 2024).

Our objective was to test whether the lack of ‘across-generation sympatric reinforcement’ influences the strength of ‘within-generation haploid-diploid reinforcement’ (Figure 1) as a barrier to hybridization. We hypothesize that 1) if conspecific sperm precedence exists in salmon/trout, its strength is influenced by ‘across-generation sympatric reinforcement’ (Figure 1b). In Europe, sympatric populations are reported to have a strong ability to enact conspecific sperm precedence (Yeates et al. 2013). We predict that 1A) due to being historically allopatric, the ability of female North American salmon to exert conspecific sperm precedence against trout is weak, however, 1B) since introduced brown trout have only been separated from Atlantic salmon for limited generations female trout will retain a strong ability to exert conspecific sperm precedence against salmon. We also hypothesize that 2) if conspecific sperm precedence exists in Newfoundland salmon/trout, then based on work from Europe it is probable that ovarian fluid is important in enabling it. If this is true, then we predict 2A) that if the ovarian fluid associated with a group of eggs is switched to the other species’ ovarian fluid, the portion of hybrids produced in sperm competition will increase, and 2B) if all ovarian fluid is removed, we predict that the male with higher quality sperm will win the sperm competition regardless of species.

Our study is the first to examine the strength of conspecific sperm precedence in North American Atlantic salmon, and examine the link between sympatric and haploid-diploid reinforcement. As salmon stocks decline in the North Atlantic (Dadswell et al., 2022), our research can be imperative in revealing a novel issue that could further threaten this species.

## Methods

### Fertilization design

With a split-clutch and split-ejaculate design, the experiment used fertilization in isolation and in sperm competition, alongside sperm swimming performance analyses to test the ability of females to exert conspecific sperm precedence under different conditions. Replication was achieved using randomized blocks on different days with unique fish (one male and one female of each species) in each block (Figure 2).

**Figure 2:**
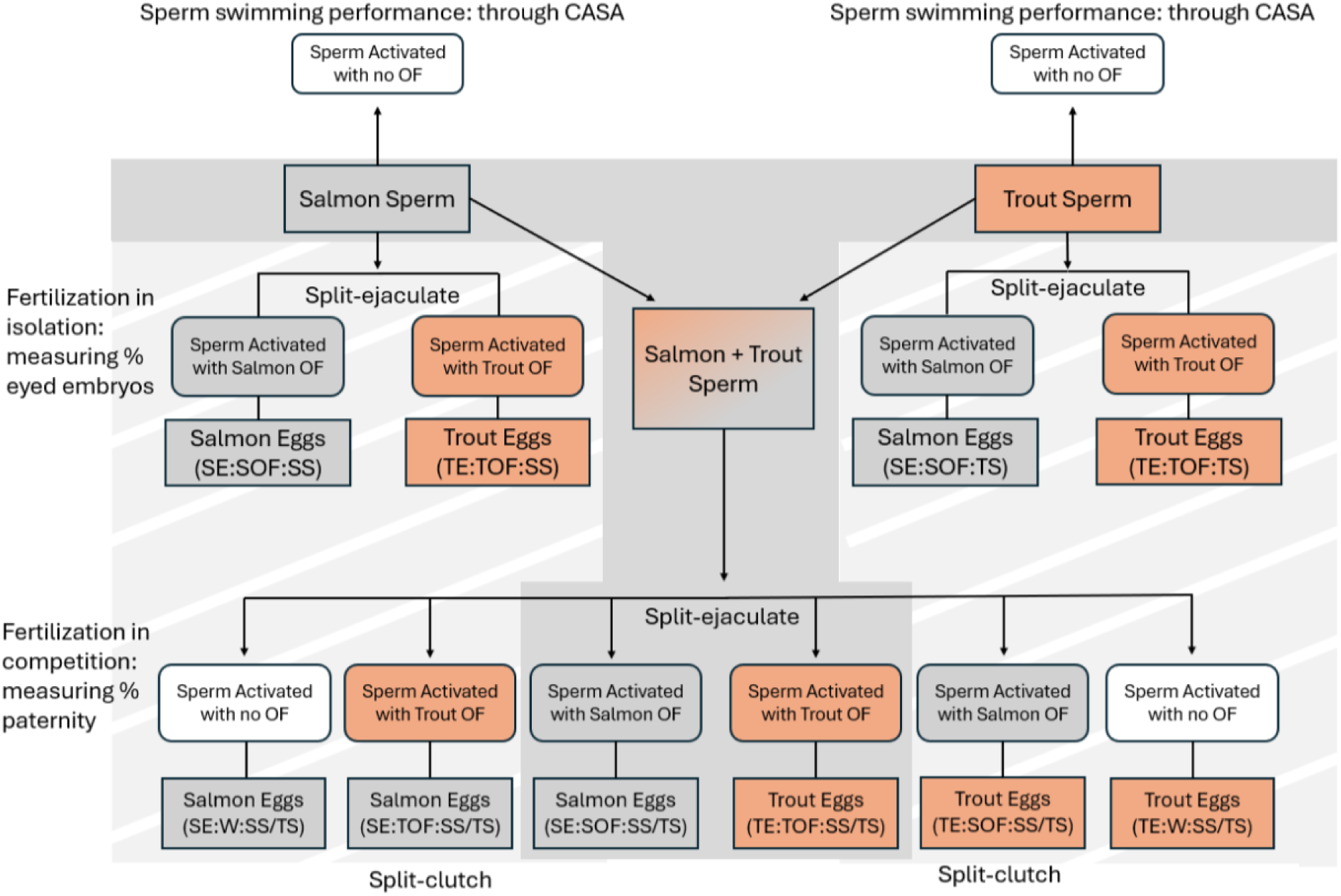
Conceptual design of each individual (n = 9) full block (all areas shaded in grey). In-vitro fertilizations were conducted for salmon and trout males in isolation and sperm competition, using one female and male of each species. Using a split-ejaculate and split-clutch technique the effectiveness of each female’s ability to exert conspecific sperm precedence was tested in three contexts related to the presence of ovarian fluid (OF) during gamete interactions: conspecific ovarian fluid (her own), the heterospecific species’ ovarian fluid, and with no ovarian fluid. Seven of the nine full blocks included sperm swimming data (CASA), and an additional (n = 6) partial blocks (areas only shaded in dark grey) provided extra unique fish to test (Hypo 1) the outcome of sperm competitions involving treatments SE:SOF:SS/TS and TE:TOF:SS/TS.

Details are described in subsequent sections. In short, both females (salmon and trout) within each block were first fertilized by each male (salmon and trout) within a block under isolation to assess gamete viability and compatibility between species and ensure that all males had the ability to fertilize eggs when sperm competition was not occurring. Then, using the same males (Figure 2) under sperm competition (SS/TS: salmon sperm / trout sperm), to test predictions 1A and 1B (the natural state) we used salmon eggs with salmon ovarian fluid (SE:SOF:SS/TS; conspecific to the eggs) and trout eggs with trout ovarian fluid (TE:TOF:SS/TS). For prediction 2A, ovarian fluid was swapped so that salmon eggs were fertilized in the presence of trout ovarian fluid (SE:TOF:SS/TS; heterospecific to the eggs) and vice versa (TE:SOF:SS/TS), and for prediction 2B eggs were fertilized without any ovarian fluid in just water (SE:W:SS/TS, TE:W:SS/TS). Sperm swimming performance analyses were performed to see how sperm quality affected the proportion of paternity a male received when competing in the absence of ovarian fluid.

In total 17 blocks were completed (Appendix 1), but due to limited embryo availability in two, nine full blocks and six partial blocks (30 males & 30 females) were used in analyses. Partial blocks (Figure 2) only contained sperm competition conditions in the natural female state to test predictions 1A and 1B.

### Study populations

North American Atlantic salmon are genetically differentiated from European populations, including in numbers of chromosomes and high levels of genome-wide divergence (Brenna-Hansen et al., 2012; Lehnert et al., 2020; Lubieniecki et al., 2010). The Atlantic salmon used in this experiment were collected in 2023 from the Exploits River in central Newfoundland, Canada (48.93 N, 55.67 W), which has not been invaded by brown trout (MacDonald et al., 2024) and thus have been allopatric from brown trout for over 600,000 years (King et al., 2007; Lehnert et al., 2020). The brown trout were taken from Windsor Lake, St. John’s, Newfoundland, Canada (47.6 N, 52.77 W), which were introduced in 1883 from Loch Leven Scotland (Hustins 2007). That population was separated from salmon when sluice gates were installed on the loch in 1831 (May, 2018). There are no salmon in Windsor Lake, and therefore the brown trout there had not been in contact with Atlantic salmon for ∼192 years, which is 38-64 generations (based on a generation time of 3-5 years).

### Fish capture and gamete collection

Salmon were captured from the Exploits River fish ladder on the Grand Falls in late September and were held in flow-through tanks at Environment and Resource Management Association (ERMA) facilities until spawning took place in November (similar to Lantiegne & Purchase, 2023; Purchase & Rooke, 2020). Trout were caught using dipnets in the tributaries of Windsor Lake during the spawning season. Through considerable coordination, within a block, gametes were collected (Nov 5-19, 2023) from both salmon and trout at the same time (<1 hour difference). After drying the abdomen with paper towels, gametes were stripped by gently squeezing the abdomen towards the vent (Purchase & Rooke, 2020). Semen samples from each fish were kept in individual plastic tubes while the eggs and ovarian fluid form each female were kept in individual insulated glass jars. All containers were kept on ice until they reached the laboratory, where they were kept in a temperature-controlled room at 4°C. Experiments took place approximately 12 hours after sampling at ∼ 10pm.

### Gamete preparation

Two teams of people worked on female and male gametes at the same time. Ovarian fluid was separated from each clutch of eggs by gravity using an aquarium net and kept in a glass beaker ∼10 minutes prior to fertilizations. For each treatment, one-part ovarian fluid and two-parts water were mixed together (33% concentration). We chose a relatively high concentration of ovarian fluid to ensure that any effects of ovarian fluid were pronounced and thus detectable. This was similar to the concentration used in other studies on sperm swimming performance (Diogo et al., 2010; Lantiegne & Purchase, 2023; Urbach et al., 2005). Ovarian fluid / water mixtures were prepared minutes before fertilizations and were kept in a temperature-controlled room at 8°C.

The eggs were washed with a 9 ppt saltwater solution to remove any remnant ovarian fluid (Lantiegne et al., 2025). Egg sizes were measured for a subset of the fish and were similar for salmon (5.84cm for 10 eggs in a straight line, on average across 7 fish) and trout (5.89cm, n = 7 fish). For each female, eggs were measured in a graduated cylinder to obtain 13mL (∼100 eggs) for fertilization under isolation and 19mL (∼150 eggs) for each fertilization under sperm competition. These measurements were acquired by individually counting 100 and 150 eggs from the female trout in our first block and measuring them. Each clutch of eggs was split into five groups (Figure 2): for salmon females this was salmon eggs in salmon ovarian fluid fertilized by salmon sperm (SE:SOF:SS) and trout sperm (SE:SOF:TS) for isolations, and salmon eggs fertilized by both salmon and trout sperm with salmon ovarian fluid (SE:SOF:SS/TS), trout ovarian fluid (SE:TOF:SS/TS) and no ovarian fluid (SE:W:SS/TS) for sperm competitions. Trout eggs followed the same organization (TE:TOF:SS/TS; TE:SOF:SS/TS; TE:W:SS/TS). Eggs were prepared minutes before fertilizations and were kept in a temperature-controlled room at 8°C.

Subsamples of both male’s semen were activated in water and checked under a microscope to ensure they were motile. Another subsample from each male was centrifuged in hematocrit tubes to measure the spermatocrit as a proxy for sperm density within the semen (Lantiegne & Purchase, 2023), so that sperm numbers could be matched between the competing males. To do this, a set volume of 126.6µL of trout (T) semen was used for each isolation treatment while of 95µL of trout (T) semen was used in sperm competition. Salmon (S) semen volumes for both isolations and sperm competition were adjusted based on the spermatocrits of both species (Equation 1). The average sperm size is similar between Atlantic salmon and brown trout (Fenkes et al., 2019; Gage et al., 2002).

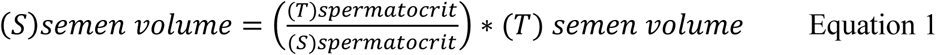

### In-vitro fertilizations

After gametes were prepared, in-vitro fertilizations for both isolation and competition groups were performed at the same time. For treatments under isolation, semen from each male was pipetted into the associated ovarian fluid solutions (SE:SOF:SS, SE:SOF:TS; TE:TOF:SS, TE:TOF:TS), which activated sperm motility. Immediately after, this was poured onto the appropriate eggs, the beaker was repeatedly swirled to enable mixing and then left to sit for three minutes (Yeates et al., 2013). For treatments under sperm competition, semen from both fathers were mixed into a 1.7mL Eppendorf tube (Figure 2). Within a few seconds, mixed semen was pipetted directly into the ovarian fluid solutions or water (SE:SOF:SS/TS, TE:TOF:SS/TS; SE:TOF:SS/TS, TE:SOF:SS/TS; SE:W:SS/TS, TE:W:SS/TS), and the same process followed.

The fertilized eggs were then rinsed using clean water through an aquarium net for ∼30 seconds. The egg beakers were filled with water and left in the dark overnight in a temperature-controlled room at 8°C. After 8-10 hours, eggs were disinfected with a 1% iodine solution (PVP iodine/ovadine) for five minutes (Fry et al., 2015). Any eggs that had turned white were deemed unviable (i.e. could not be fertilized) and were discarded (Lantiegne et al., 2025).

For each treatment under isolation, 100 viable eggs were placed as batches of 50 in PVC tubes (5.8cm height x 5.8cm diameter) with a screen on the bottom. For treatments under sperm competition three tubes of 50 eggs were used in the same way. Tubes were placed inside of one of two MariSource^TM^ 4-tray vertical flow-through incubators, which were located inside separate environmental chambers that were temperature controlled at 8°C, alternating by block.

Incubators used a recirculating system where water was running over the eggs. Salmonids are sensitive in their early embryonic stage (Musialak et al., 2024), and can die when exposed to light or mechanical stimuli. Therefore, incubators were kept in a dark room where physical contact was minimized. While incubating, embryos were kept in the dark, and only red lighting was used when checking for mortality. Water was continually recycled and treated via mechanical and UV filtration, and 23 liters of water were added every other day to account for evaporation.

### Data collection

#### Semen quality

Subsamples of semen from each male were assessed for sperm swimming performance (Figure 2). Using a separate aliquot, seminal fluid was separated from sperm by centrifugation, and was re-added to the test subsample at a 75:1 ratio with semen to dilute the number of sperm cells on a slide (Lantiegne & Purchase, 2023; Purchase & Moreau, 2012). Bovine serum albumin was added to the semen and seminal fluid mixture at a 1:1000 concentration to prevent the sample from sticking to the microscope slide (Beirão et al., 2014; Lantiegne & Purchase, 2023). Subsamples awaiting the microscope were kept cool using a Torrey Pines Scientific Echotherm set at 5°C and when placed on the microscope slide were cooled using a custom Physitemp TS-4 system set at 9°C.

Each male’s sperm was recorded in a swimming medium of a 1:1 concentration of either salmon or trout ovarian fluid with water or with just water itself. We used a 50% concentration of ovarian fluid (Rosengrave et al., 2008, 2009) as this has been found to maximize sperm swimming speed (Turner & Montgomerie, 2002), which should exaggerate any differential effect sizes between SOF and TOF if they existed. Sperm swimming videos were recorded using Streampix software, on a Prosilica GE680 camera attached to an inverted Leica DM IL LED microscope. Videos were checked to ensure that video quality was acceptable and that a sufficient number of sperm cells (40-200) were visible. Three technical replicates were taken for each combination of sperm type and swimming medium (SOF:SS, SOF:TS, TOF:SS, TOF:TS, W:SS, W:TS) for a total of 18 videos per block. Sperm movement videos were then analyzed using CASA_automated (Purchase & Earle, 2012), a modified version of the Computer Assisted Sperm Analysis (CASA) plugin for ImageJ (Wilson-Leedy & Ingermann, 2007). Salmonid sperm have short lifespans, and fertilize rapidly (Hoysak & Liley, 2001). Therefore, sperm were tracked at 6-7 and 7-8 seconds post-activation (and then averaged between the two time periods), the earliest at which sperm activity could be reliably recorded. Three qualities of sperm swimming performance were measured: sperm motility (MOT), and the curvilinear velocity (VCL) and linearity (LIN) of motile cells, which are important in sperm competition (Boschetto et al., 2011; Gage et al., 2004), for each male when either salmon, trout, or no ovarian fluid was present. These were then compared with the same male’s paternity rates in fertilization treatments under sperm competition with no ovarian fluid.

#### Male success when in isolation from the other male

The 100 viable eggs exposed to sperm for single males under isolation for each treatment (i.e. one male + one female) and subsequently incubated were counted as either successful or unsuccessful in developing embryos that reached the eyed stage (combination of fertilization success and survival).

#### Paternity assignment under sperm competition

The mother of each embryo was known; however, paternity was unknown when salmon and trout males were under sperm competition. To determine the outcome of sperm competition, preserved eyed embryos were used for genetic analyses to identify the species of their father.

Embryos were haphazardly selected from each treatment for paternity testing. DNA from each embryo was extracted from 1mm^2^ fin clips in a 96-well plate using a Chelex-based procedure taken from Elliott et al., (2020). These samples were then used in PCR, using a reaction that included 12.5µL of DreamTaq Green PCR MasterMix (2X), 1µL of primers (A 5’TACGCCCGATCTCGTCCGATC-3’) and (B 5’CAGGCTGGTATGGCCGTAAGC-3’; Pendas et al., 1995), 8.5µL of RNAse free water, and 2µL of an individual embryo’s DNA. Reactions were placed into a thermal cycler at 95°C for five minutes, then put on a cycle going from 94°C - 60°C - 72°C for 30 seconds each and repeated for 35 cycles, then a final temperature of 72°C for five minutes to complete the process. Within 24 hours of PCR, samples were run using gel electrophoresis with a 2% agarose solution which contained 7µL of SYBR safe DNA gel stain. Gels were run at either 75V for two hours or 120V for one hour and scanned using Bio-Rad ChemiDoc imaging system, where the species of each embryo could be determined depending on the size and number of bands (Appendix 2).

Positive controls were created with known salmon, trout, and hybrid DNA, which were taken from individuals created in isolated in-vitro fertilizations. These were used alongside unknown samples in sperm competition within each gel to verify the species of embryos and the integrity of paternity assignment. Embryos were genotyped in batches of 6 per treatment, and the total number (n = 18-48) genotyped in a treatment depended on how dominant one male was compared to the other. Fewer embryos were added to the 18 minimum if one male secured a vast majority of paternity. This was tested in several groups where additional embryos did not change outcomes. A total of 1878 embryos were genotyped (Appendix 3). After genotyping embryos in sperm competition, the number of pure and hybrid embryos were scored across each female species within each treatment.

#### Paternity standardization to test for conspecific sperm precedence

To evaluate conspecific sperm precedence, we standardized paternity rates between salmon-trout males in sperm competition using the same male’s performance in isolation. Each embryo assigned paternity to a given male under sperm competition represented a combination of egg fertilization and embryo survival to that development stage. Variation in either process would influence relative perceived paternity, but only female alteration of fertilization is indicative of conspecific sperm precedence. The two competing males within a block could have different sperm qualities (e.g., based on their diet), which could affect fertilization success without female influence. Hybrid embryos may have a lower survival rate to the eyed stage, and thus paternity of survivors might be biased against hybrids even if hybrid fertilizations were very high. For example, if the conspecific male did better than the heterospecific male in isolation, then the conspecific male would be assigned higher paternity under sperm competition, even if the female did not make any modifications to fertilization. This can create noise that would have an impact on the ability to assess whether or not conspecific sperm precedence is occurring.

To account for this, we used each individual male’s proportion of eggs that fertilized and survived in isolation to add a weight to performance in competition. By taking the number of embryos fathered by a male salmon (s1) and trout (t1) in sperm competitions and the success each male (Ps, Pt) got in isolation (with the same female), our model extrapolated the data to output the number of total embryos belonging to each father (s0, t0) in sperm competition out of all 150 viable eggs (T) used in each competition, rounded to the nearest whole number. A hypothetical example follows. When fertilizing a group of salmon eggs, the male salmon could have gotten 100% embryo development in isolation and the male trout got 75%. In sperm competition between the same males over the same female, the salmon and trout male might have fathered 48/96 embryos each (50% paternity), meaning the male trout received a boost. To better visualize the effects of conspecific sperm precedence, our model standardized our sperm competition values as if performance in isolation between males were equal:

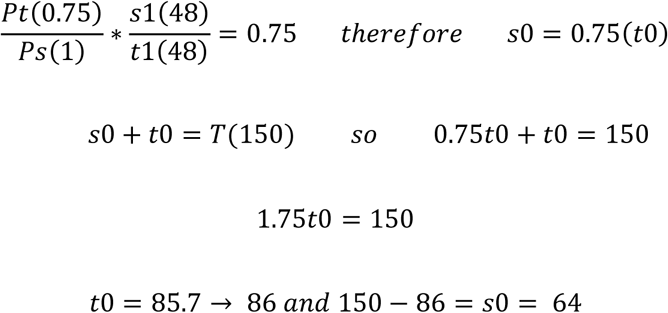

To perform such standardization, it was important to record embryo success in isolation and sperm competition at the same development stage, as if fertilization success in isolation was measured earlier than tissue was available for paternity, differential mortality would impact the result. Salmonids produce large eggs and have slow development under cold water conditions. Paternity is usually performed once embryos develop enough such that their eyes are observable. Eyes were developed in embryos when sampled at 264 ATUs (accumulated temperature units), which was 33 days post-fertilization for each block. Eyed and non-eyed eggs were counted in isolation (Appendix 1), and eyed eggs from sperm competitions were preserved with 95% ethanol in separate tubes by treatment by block.

Atlantic salmon embryos reach the feeding stage after the yolk sac is completely absorbed at “emergence” in about 900 ATUs (Gorodilov, 1996). We therefore measured the paternity of our embryos ∼29% of the way through development to the free-living stage. This is relatively early in development (and thus less confounding variables from differential mortality) compared to commonly studied species such as zebrafish, in which paternity is usually identified around 40-60% of the way through development (Coe et al., 2009; Watt et al., 2011), or fruit flies, which are not identified until the start of the feeding stage (Harshman & Clark, 1998; Hurtado et al., 2013).

### Data analyses

#### Male success in isolation without sperm competition

We evaluated the compatibility of Atlantic salmon and brown trout by examining the difference in the fertilization and survival of resulting embryos in both conspecific and heterospecific breeding pairs. If heterospecific pairs were to produce less eyed embryos than conspecific pairs, conspecific males would be perceived biased towards in competition without conspecific sperm precedence necessarily ever occurring. We compared the successes and failures between each combination of breeding pair using a generalized linear mixed model with a binomial distribution (Equation 2) for 9 blocks (18 males, 18 females) of all four of our isolation treatments (SE:SOF:SS, SE:SOF:TS; TE:TOF:SS, TE:TOF:TS). The response variable was the # of eggs that were fertilized and survived to the eyed stage (successes) and the # of eggs that did not develop to the eyed stage (failures). Our fixed factors were the species of the male, the species of the female, and their interaction, while block was used as a random factor. The objective of this test was to see if the interaction between male and female species was significant in determining the # of successes and failures. As the interaction was significant, we created a post hoc test that used pairwise comparisons between treatments to determine whether the eyed embryos produced by conspecific males was significantly higher than those produced by heterospecific males within both our female salmon treatments (i.e. SE:SOF:SS vs SE:SOF:TS) and female trout treatments (i.e. TE:TOF:TS vs TE:TOF:SS).

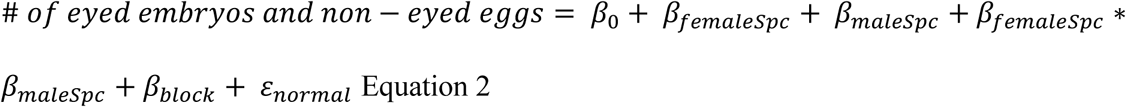

#### Sperm competition between males in the natural ovarian fluid state

Predictions for hypothesis #1 were that Atlantic salmon would have a weak ability to exert conspecific sperm precedence while brown trout would have a strong ability to do so. We tested this in the natural state, where both Atlantic salmon and brown trout eggs were fertilized within their own ovarian fluid. We used a generalized linear mixed model with binomial distribution (Equation 3) to evaluate our predictions. The response variable was the # of purebred embryos and hybrid embryos in associated sperm competition treatments (i.e. SE:SOF:SS/TS vs. TE:TOF:SS/TS), while the species of the female was included as a fixed factor and block as a random factor. No interaction was included as only the species of female were being compared.

Our generalized linear mixed model was performed three times using three different datasets. We compared raw paternity results that were not adjusted for male success in isolation for both all 15 blocks (30 males; 30 females), and subsequently for the nine blocks (18 males; 18 females) that were used in the standardization process (Figure 2, Appendix 1), with an added zero inflation component (which improved model diagnostics) to account for many competitions that were completely dominated by one male. Then we reran the analyses again for our standardized sperm competition values, creating another generalized linear mixed model for our nine full blocks with no zero inflation component.

Using the same model, we also evaluated for the strength of conspecific sperm precedence in each female species. To do this we compared the # of pure and hybrid embryos in our standardized dataset (to account for performance in isolation) within either salmon females or trout females to a simulated dataset where the # of pure and hybrid embryos was equal using the “emmeans” package (Lenth & Piaskowski, 2023). This allowed us to see if there was a significant difference in our results when compared to a scenario where no conspecific sperm precedence is occurring.

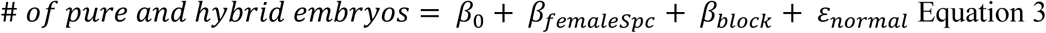

#### Effects of ovarian fluid on conspecific sperm precedence

Prediction 2A was that if the conspecific ovarian fluid associated with a group of eggs is artificially swapped to be heterospecific, the proportion of hybridization increases. We tested this by evaluating if the # of pure and hybrid embryos changed between treatments (Figure 2) using a zero-inflated generalized linear mixed model with binomial distribution (Equation 4) on nine blocks (18 males; 18 females), which included all four treatments in sperm competition that contained ovarian fluid (SE:SOF:SS/TS; SE:TOF:SS/TS; TE:TOF:SS/TS; TE:SOF:SS/TS). The response variable was the # of pure and hybrid embryos, and our fixed factors were species identity of the eggs and ovarian fluid (either SOF or TOF) and their interaction, while block was used as a random factor. As the interaction between ovarian fluid and the species of the eggs was significant, post hoc pairwise comparisons were performed to see if there was a significant difference in the # of hybrid embryos within each female’s eggs (i.e. SE:SOF:SS/TS vs SE:TOF:SS/TS; TE:TOF:SS/TS vs TE:SOF:SS/TS).

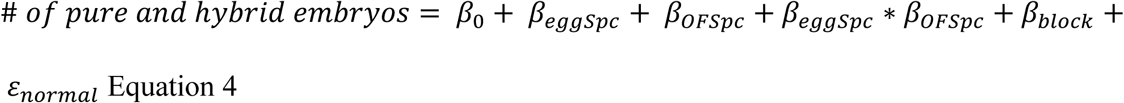

All models were fitted in R version 4.5.2 (R Core Team, 2025). Generalized linear mixed models were fitted using the “glmmTMB” package (Brooks et al., 2017). Model diagnostics were performed with the “DHARMa” package (Florian, 2016), and post-hoc analyses were conducted using the “emmeans” package (Lenth & Piaskowski, 2023) with proportional weighting (weights=“prop”).

For prediction 2B, we predicted that if no ovarian fluid is present, sperm quality decides which individual wins a higher portion of fertilization. We fitted linear mixed effects models for our treatments that did not contain ovarian fluid (SE:W:SS/TS; TE:W:SS/TS) in the seven blocks (14 males; 14 females) that had sperm swimming performance data (Figure 2). The response variable was the % difference in paternity between males, with our fixed factors being either the % difference in MOT, VCL, or LIN between males, as well as egg species and their interaction, and with block as a random factor (Equations 5-7). The values used in our models for the three sperm performance qualities were averaged across all three swimming mediums (SOF, TOF, W) to create a robust indicator for semen quality differences between males, but were also performed in each medium individually (Appendix 4). No interactions were significant and thus no post hoc tests took place. Residuals were checked using Q-Q plots and normality was observed in all models. All linear mixed effects models were fitted using the “lme4” package (Bates, 2007).

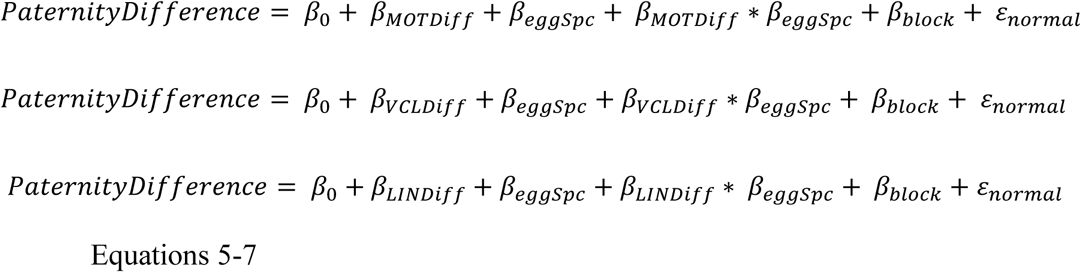

## Results

### Male success in isolation without sperm competition

Of the 2000 viable eggs incubated from each species under male isolation, 1038 salmon eggs and 1326 trout eggs were fertilized and survived to the eyed stage. We found that the interaction between male and female species was significant (χ² = 86.9913, df = 1, p < 0.0001), meaning that the # of embryos that were fertilized and survived to the eyed stage was different across breeding pairs. Our post hoc testing revealed that in salmon females, salmon males produced significantly more offspring (combination of fertilization and survival to eyed embryo stage) than trout males (SE:SOF:SS, z = 6.174, p <0.0001), while in trout females, trout males produced more offspring than salmon males (TE:TOF:TS, z = −7.011, p < 0.0001). Averaged across females within a species (Figure 3), the percentage of eggs that were fertilized and survived to the eyed stage was 21.4% lower for hybridized salmon eggs (50.8%) compared to pure salmon (64.6%), and 17.6% lower for hybridized trout eggs (66.6%) compared to pure trout (80.8%). Our results indicate that salmon and trout gametes were compatible, and that conspecific pairs can produce more eyed embryos through either fertilization or survival, which should be accounted for in sperm competition comparisons.

**Figure 3:**
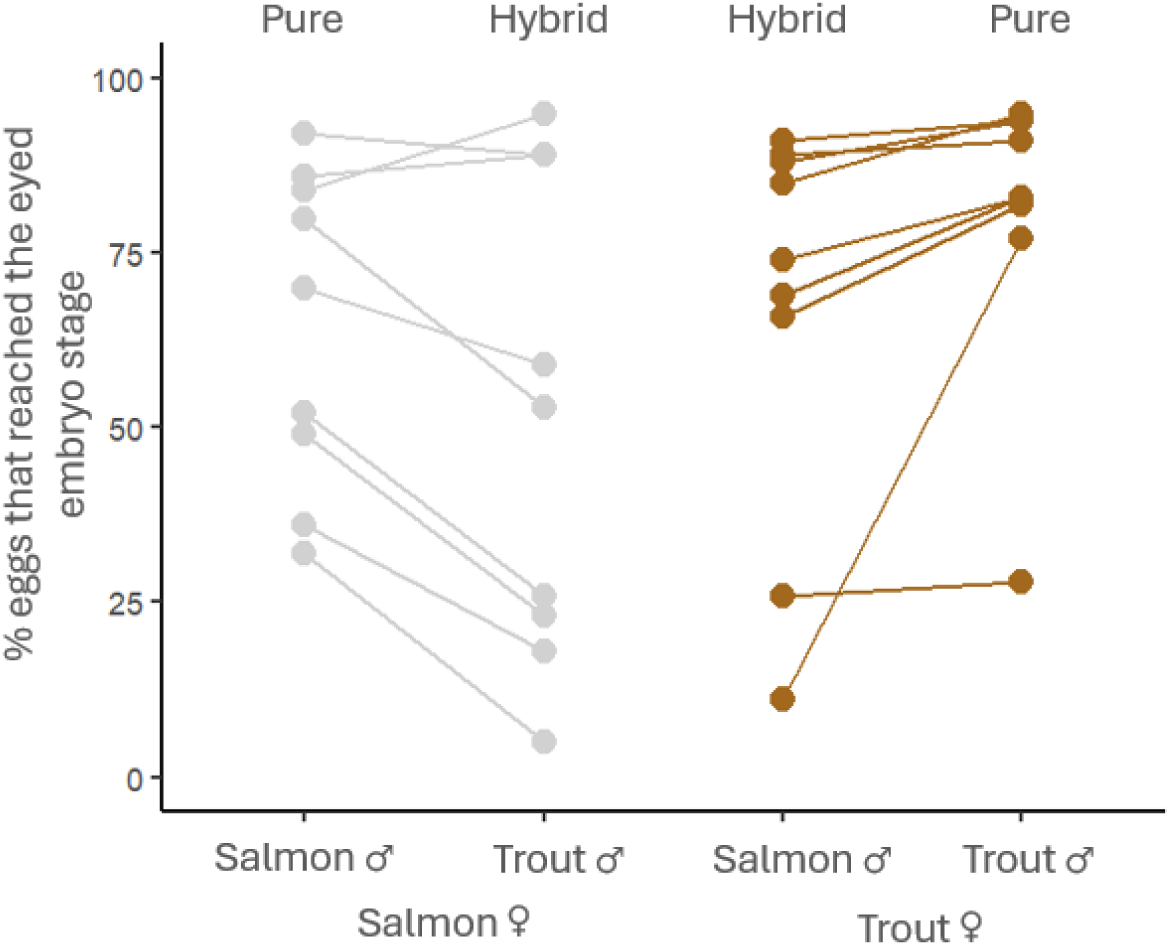
Percentages of embryos that reached the eyed stage for each female’s eggs when fertilized by a male trout or salmon in isolation (no sperm competition). Lines represent each female within the block design, and the species of female is represented by color. The same males were used between species of female within a block. Across females, on average, hybridized eggs had 21.3% and 17.6% fewer eyed embryos than pure sibling eggs for salmon and trout females, respectively. The reduction was not dramatic nor universal and stemmed from an unquantified combination of different fertilization success and survival of fertilized eggs to the eyed stage.

### Sperm competition between males in the natural ovarian fluid state

When having her eggs fertilized in the natural ovarian fluid state, we found that female salmon did not exert conspecific sperm precedence while female trout had a strong ability to do so, which supports predictions 1A and 1B. We found a significant difference in the # of purebred and hybrid embryos between species in our natural ovarian fluid treatments (Figure 4a, n = 15 blocks, χ² = 13.0500, df = 1, p = 0.0003; Figure 4b+c, n = 9 unstandardized blocks, χ² = 9.3118, df = 1, p = 0.0023; Figure 4b+c, n = 9 standardized blocks, χ² = 68.6235, df = 1, p < 0.0001) which indicates that one species produced more conspecific offspring than the other. In our standardized competition dataset conspecific males did not achieve significantly higher paternity in female salmon eggs (50.6% salmon paternity, 49.4% trout paternity; z = 0.011, p = 0.9915, Figure 4b), but did in female trout eggs (68.0% trout, 32.0% salmon; z = 3.387, p = 0.0007, Figure 4c). Standardizing paternity in sperm competition using individual male performance in isolation creates a clearer comparison of female influence and accounted for the known significant difference in fertilization and/or embryo survival between conspecific breeding pairs.

**Figure 4:**
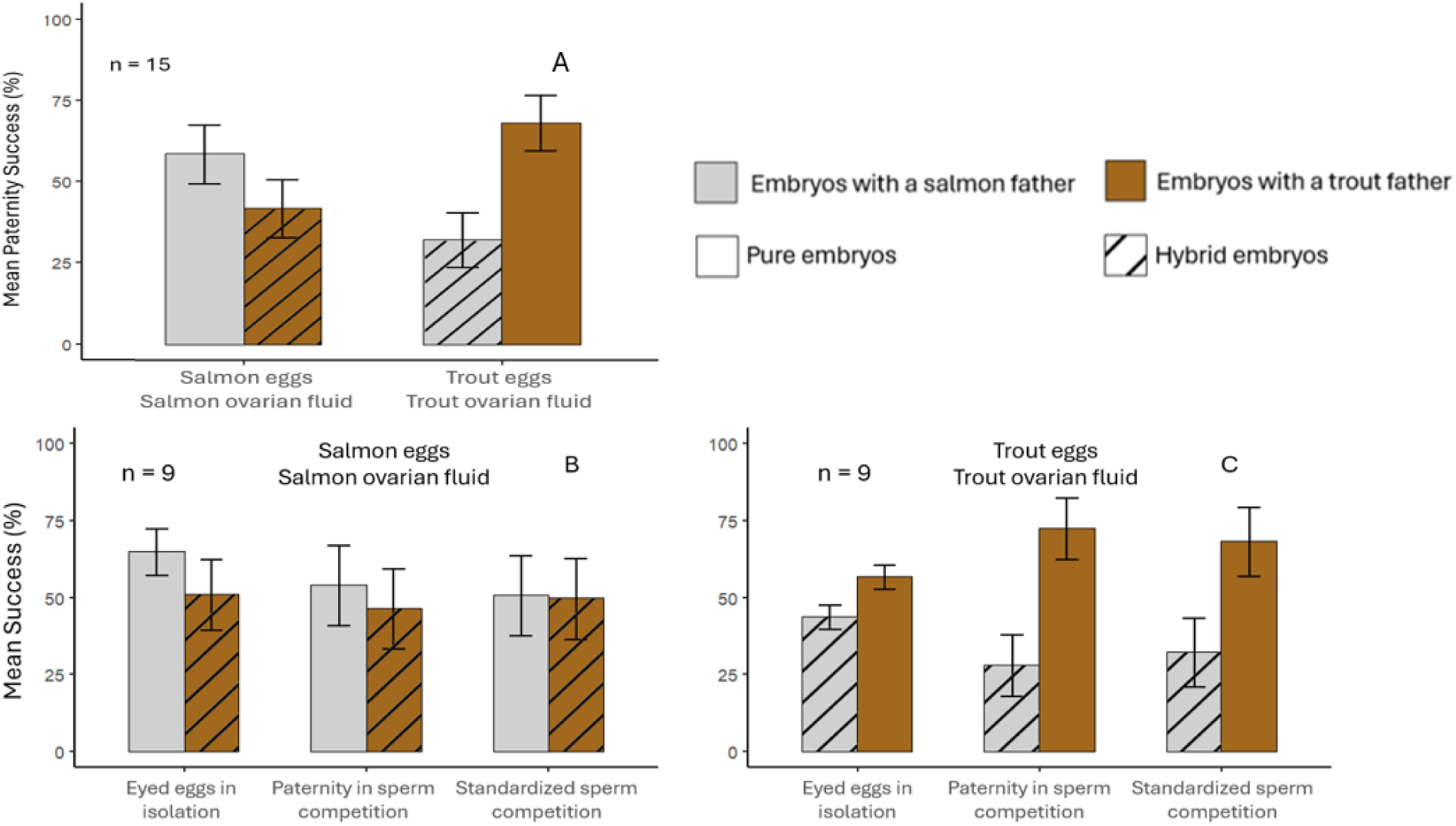
A) Mean paternity success across blocks between salmon-trout males under sperm competition in the natural egg state (SE:SOF, TE:TOF) for both species of female in 15 blocks (60 adult fish). B&C) Mean success between salmon-trout males under isolation and sperm competition in the natural state (e.g., SE:SOF:SS, SE:SOF:TS, SE:SOF:SS/TS), showing unstandardized and standardized values for both species of female in 9 blocks (36 adult fish). Hybrid embryos are represented by hatched bars. Relative paternity between males in both sperm competition and standardized groups sum to 100. Error bars represent standard error.

### Effects of ovarian fluid on conspecific sperm precedence

The interaction between ovarian fluid and eggs was significant (χ² = 30.01, df = 1, p < 0.0001) in determining the # of pure and hybrid embryos between treatments. When ovarian fluid accompanying a female’s eggs was replaced with heterospecific ovarian fluid (Figure 5), the number of hybrids produced under sperm competition between male trout and salmon increased significantly (prediction 2a) in both species (SE:TOF, z = 3.180, p = 0.0080; TE:SOF, z = −4.674, p <0.0001), supporting hypothesis 2. In salmon eggs, salmon males achieved 53.7% of paternity on average when conspecific SOF was present, but the same individuals got 37.7% when it was swapped for heterospecific ovarian fluid. In trout eggs, trout males achieved 72.1% paternity when conspecific TOF was present, but 52.6% with SOF. On average, hybridization increased by 42.3% (salmon females 25.7%, trout females 58.9%) when ovarian fluid was swapped.

**Figure 5:**
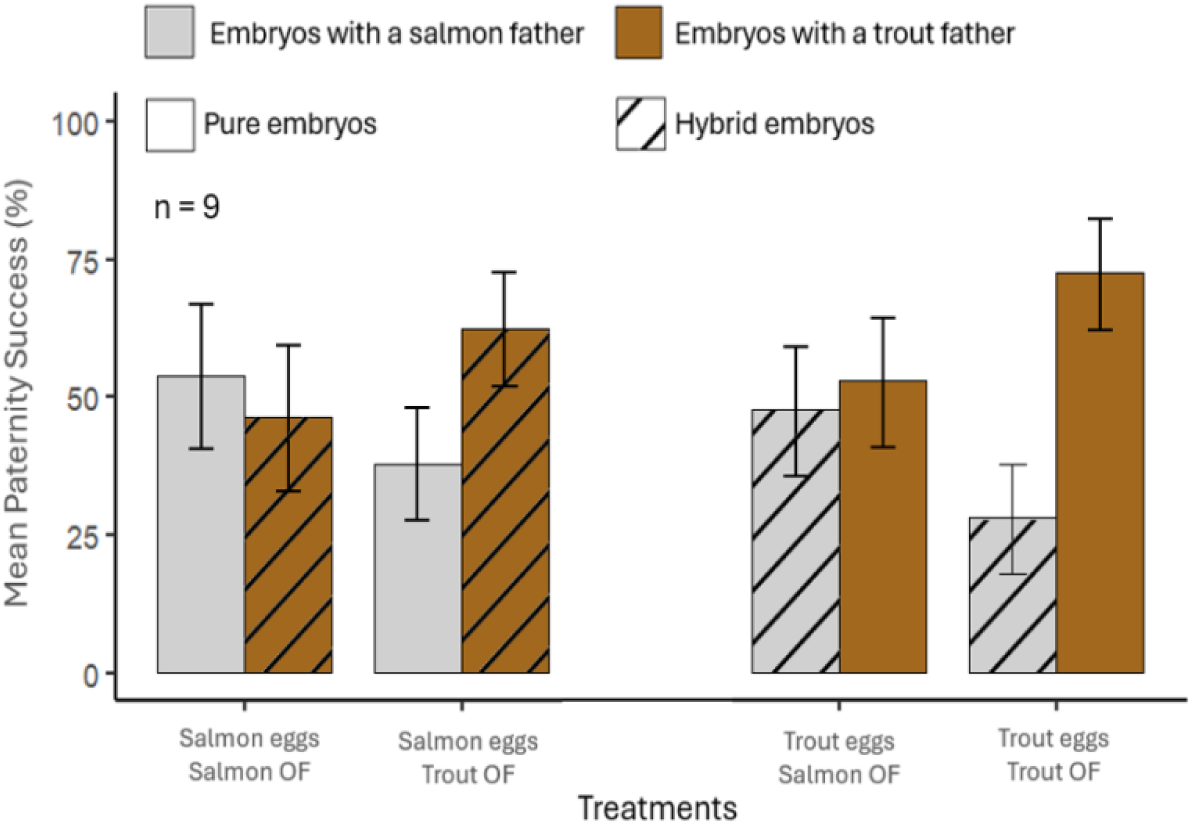
Mean paternity success across blocks between salmon-trout males under sperm competition for both the natural state (SE:SOF:SS/TS, TE:TOF:SS/TS) and swapped ovarian fluid groups (SE:TOF:SS/TS, TE:SOF:SS/TS) for both maternal species. Hybrid embryos are represented by hatched bars. Relative paternity between males in each group sums to 100. Error bars represent standard error among 9 blocks (36 adult fish). Hybridization rate of a female’s eggs increases if conspecific ovarian fluid is replaced with that of the heterospecific species.

Using the same fish, paternity changed drastically in the absence of any ovarian fluid (unnatural situation), supporting hypothesis 2. On average, paternity was dominated by trout males (Figure 6) in both trout eggs (84.1%) and salmon eggs (75.6%), and within the majority of our blocks male trout achieved >90% of paternity. No interactions between the % difference in sperm quality and the species of the egg were found to be significant in determining the difference in paternity (% sperm that were motile: χ² = 0.1027, df = 1, p = 0.7486; curvilinear swimming velocity of motile cells; χ² = 0.5901, df = 1, p = 0.4424; swimming linearity of motile cells; χ² = 0.0565, df = 1, p = 0.8121). Contrary to prediction 2B, within a block, the relative semen quality of each male (Figure 6, Appendix 4) did not clearly explain relative paternity (% sperm that were motile: χ² = 2.41, df = 1, p = 0.1206; and curvilinear swimming velocity of motile cells; χ² = 0.61, df = 1, p = 0.4356; swimming linearity of motile cells; χ² = 1.67, df = 1, p = 0.1969).

**Figure 6:**
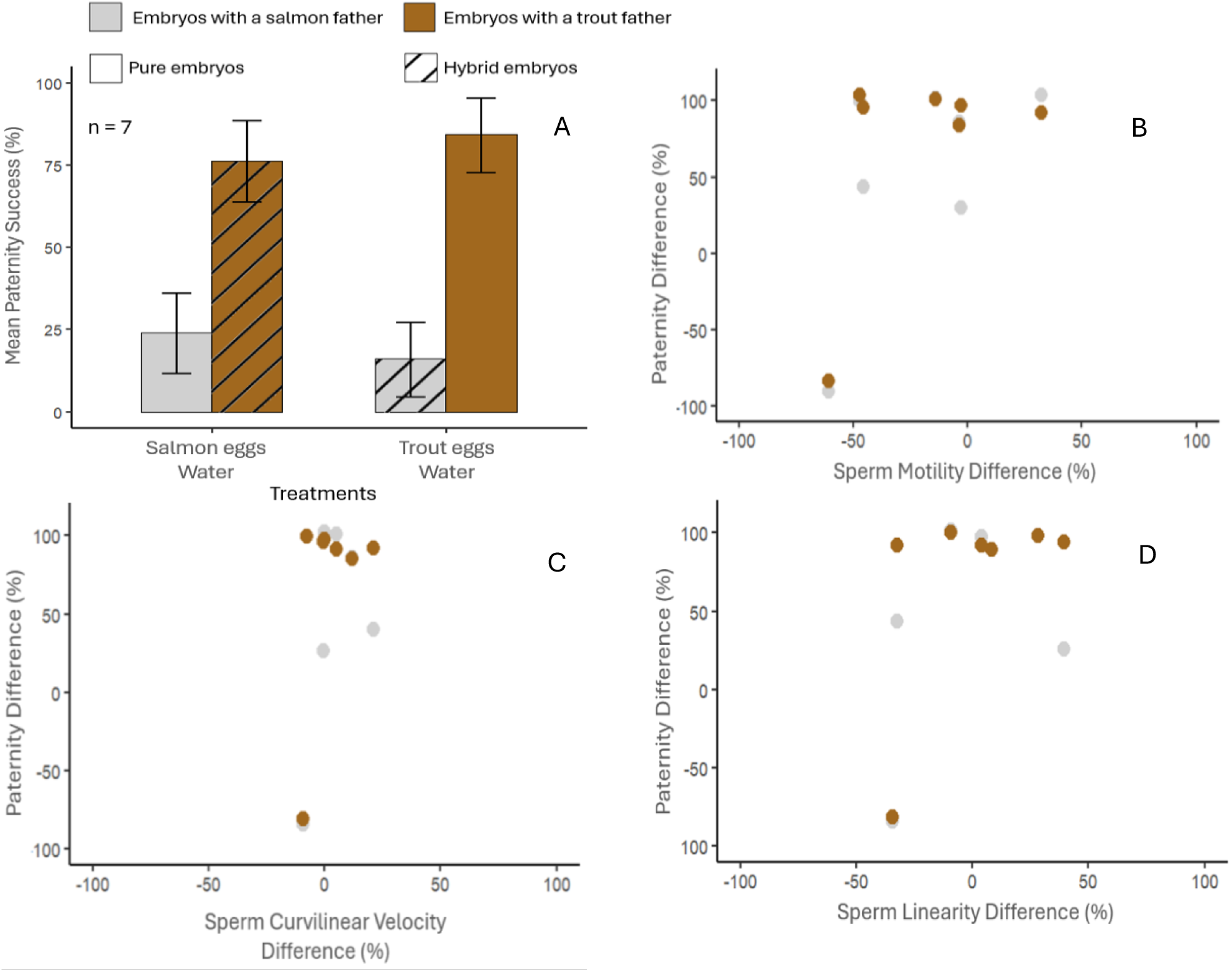
A) Mean paternity success across blocks between salmon-trout males under sperm competition without the presence of ovarian fluid (SE:W:SS/TS, TE:W:SS/TS) in both species of female. Hybrid embryos are represented by hatched bars. Relative paternity between males in each group sums to 100. Error bars represent standard error among 7 blocks (28 adult fish). Trout males dominated paternity for both trout and salmon eggs. Within a block the winner of sperm competition was not associated with sperm swimming parameters. Panels B (percent of sperm cells that were motile), C (sperm swimming velocity) and D (sperm swimming linearity) compare the difference in paternity to the difference in semen quality between trout-salmon males within each block (e.g. if trout paternity is 30% lower than salmon paternity the paternity difference is −30). The colors for panels B-D represent the species of eggs being competed over. Semen quality measurements for each male were averaged between performance in both ovarian fluids (SOF, TOF) and when no ovarian fluid was present (W).

## Discussion

We expected, and found, that when under in-vitro sperm competition Newfoundland native Atlantic salmon females could not strongly exert conspecific sperm precedence against trout, but invasive brown trout females could against salmon. Our findings contrast research in Europe, which demonstrated that sympatric populations of Atlantic salmon could exert conspecific sperm precedence to much the same extent as brown trout (Yeates et al., 2013).

Barriers to fertilization evolve rapidly, and often stem from sexual selection (Price, 1997), which is intense both within and between species of salmonids (Fleming & Gross, 1994; Garcia-Vazquez et al., 2002). Thus, in areas where two closely related salmonids live in sympatry, it would be expected that barriers to hybrid fertilizations are strong in both the pre- and post-ejaculatory selection phases. Other pairs of closely related species have demonstrated that when in sympatry, behavior in the pre-ejaculatory stage prevents hybrid matings more effectively than when the same species exist in allopatry (Gabor & Ryam, 2001; Moran et al., 2018; Moran & Fuller, 2018; Pfennig & Rice, 2014; Rundle & Schluter, 1998; Stre et al., 1997). There is a lack of information on the post-ejaculatory phase, but a study on fruit flies has indicated that the strength of sympatric reinforcement is correlated with the strength of conspecific sperm precedence (Castillo & Moyle, 2019). It seems likely that after being separated from European populations for over 600,000 years (King et al., 2007; Lehnert et al., 2020), sympatric reinforcement has weakened in North American Atlantic salmon, which has in turn, weakened their haploid-diploid reinforcement and ability to use conspecific sperm precedence. Our brown trout had only been separated from European populations for ∼192 years and our findings indicate that they still possess haploid-diploid reinforcement acquired in Europe.

Our experiment draws much inspiration from Yeates et al. (2013), however possesses some key differences, as all the fish used in our experiment were collected within the same spawning season and used consistently in both our isolation and competition groups. We also purposefully allowed our embryos in isolation to develop alongside those in sperm competition groups before scoring paternity, something that was not done in Yeates et al. (2013), to account for differential male fertilization ability and survival of his fertilized eggs when standardizing sperm competition. Our standardizations were only possible because we used the same fish and allowed embryonic development, and thus more accurately presented the extent of conspecific sperm precedence in both species, preventing the masking or over-exaggeration of conspecific sperm precedence on paternity.

No other published studies exist on the ability of North American Atlantic salmon to exert conspecific sperm precedence. Our experiment was the first to use in-vitro fertilizations between a historically allopatric Atlantic salmon and sympatric brown trout population. Due to the unique conditions of both of these populations having never interacted before, we could ensure that our results were not confounded by any contemporary sympatric reinforcement. This has helped elucidate the link between sympatric reinforcement and haploid-diploid reinforcement, and supports previous research stating that the ovarian fluid is at least partially responsible for biasing sperm competition.

To further our understanding of how these mechanisms work, more in-vitro fertilizations should be conducted using a wider range of salmonids, including more Atlantic salmon populations in Newfoundland. Certain populations of Atlantic salmon in Newfoundland have been exposed to brown trout (MacDonald et al., 2024; Westley & Fleming, 2011), and one could compare the effectiveness of conspecific sperm precedence between salmon populations. If the salmon females that co-exist with trout exhibit a stronger ability to select conspecific males, it would suggest that some reinforcement has occurred. If our predictions hold true in other experiments, the length of time in which Atlantic salmon and brown trout have been sympatric to each other will correlate with the strength of conspecific sperm precedence. Populations of Atlantic salmon and brown trout that have never been separated will have a strong ability to exert conspecific sperm precedence, but the longer a population has been isolated from the other conspecific sperm precedence will weaken. No published articles have studied post-ejaculatory, prezygotic barriers in any other pairs of salmonid species besides Atlantic salmon and brown trout, and the extent that other pairs of species would be able to use conspecific sperm precedence, if at all, is unknown. If this mechanism is ubiquitous in salmonids, then we would expect that historically sympatric populations, like for example Chinook (*Oncorhynchus tshawytscha*) and Coho salmon (*O. kisutch*), or Pink (*O. gorbuscha*) and Chum salmon (*O. keta*), which naturally hybridize (Araujo et al., 2021; Chevassus, 1979), have a strong ability to exert conspecific sperm precedence (Purchase, 2022).

Despite conspecific sperm precedence appearing to be absent in Newfoundland Atlantic salmon, hybridization increased in both species when the associated ovarian fluid was swapped. This corresponds with previous research that identifies ovarian fluid as being very important in allowing females to exert cryptic female choice in salmonids (Lehnert et al., 2017; Rosengrave et al., 2008; Yeates et al., 2013) as well as other aquatic species (Alonzo et al., 2016; Diogo et al., 2010; Geyer & Palumbi, 2005; Zadmajid et al., 2019). The exact mechanism of how salmonid ovarian fluid promotes conspecific fertilization is not understood, and current research has found mixed results around ovarian fluid’s ability to preferentially upregulate sperm swimming performance of preferred males (Lantiegne & Purchase, 2023; Rosengrave et al., 2016; Yeates et al., 2013). Regardless, when males compete when ovarian fluid is not present, we would expect females to be unable to use cryptic female choice to select the preferred male. From the data we collected we found that this was supported and there was no evidence of conspecific sperm precedence occurring, as trout dominated paternity in both females equally. It is not clear why trout were more successful in competition, as sperm performance analyses did not show any swimming qualities being significant in achieving fertilization.

Our findings suggest that Atlantic salmon females are at a higher risk of hybridization than brown trout females. While there is very little information on the direction of hybridization between these species in Newfoundland, one study found that adult hybrids were produced only by salmon fathers and trout mothers (McGowan & Davidson, 1992). While these findings were speculative at best due to a low sample size, if this was the standard across all hybrids in Newfoundland, it would seemingly contradict our findings. One explanation for this is that our experiment mimics a scenario where the number of salmon and trout are equal, but in reality, it is much more likely that in most rivers native salmon vastly outnumber invading trout. Salmon females, who are at a higher risk of hybridization in sperm competition, should be much less likely to mate with a male trout, as the number of conspecific mates should be much greater. This is opposed to trout females, which likely have far fewer options for conspecific mates, and thus may mate more often with salmon males. Another possible reason for the direction of hybridization is the presence of mature precocial salmon parr, which brown trout would inevitably encounter when invading streams already inhabited by salmon. Mature parr are indicated to increase hybridization between these species (Crozier, 1984; Jansson & Öst, 1997; Verspoor, 1988), and thus could reduce the number of pure trout offspring by stealing fertilizations from trout males, as they do in Europe (Garcia-Vazquez, 2001). As an example, competition for a female trout’s eggs may be between one trout and ten mature salmon parr, and even the ability to use conspecific sperm precedence may not be enough to prevent a majority of those eggs from being fertilized by heterospecific males. Overall, these interactions may be significant in creating a barrier that prevents brown trout from establishing self-sustaining populations in salmon dominated streams (Purchase et al., 2024).

Atlantic salmon stocks in North America have been on the decline in recent decades in part due to reduced survival rates at sea (Dadswell et al., 2022; Dempson et al., 2024; Mills et al., 2013). If the abundance of Atlantic salmon is promoting a barrier in preventing hybridization, overtime declining salmon densities could allow brown trout to invade rivers and streams inhabited by salmon more easily, and would allow female brown trout to select and produce offspring with conspecific mates more often. If more trout were present in salmon streams, more female salmon would find themselves under mating pressure from male trout, and unless those females could adapt to use conspecific sperm precedence in a relatively short amount of time, hybridization would likely increase. Hybridization may be found to be significantly lower in streams where Atlantic salmon vastly outnumber brown trout, or in streams that have a larger percentage of mature male parr. Future research should attempt to quantify the difference in the frequency of hybrids among streams with differing densities of Atlantic salmon, mature salmon parr, and brown trout.

## Acknowledgements

Funding was provided by Memorial University of Newfoundland, the Canadian Foundation for Innovation, the Natural Sciences and Engineering Research Council of Canada, and the Research and Development Corporation of Newfoundland and Labrador. We would like to especially thank the staff of the Environmental Resources Management Association for their efforts in providing the Atlantic salmon gametes used in our experiment. Niels Van Miltenburg, Madison Philipp, and Nick Murphy were of great assistance in both collecting brown trout gametes in the field and conducting in-vitro fertilizations in the lab. We also thank Ian Fleming for his comments on an earlier version of my manuscript.

## Appendices

**Appendix 1:** Left side of table shows the number of eyed embryos that could be used in paternity testing for each treatment of each block. Blocks 1-10 represent full blocks with all 5 treatments. Blocks P1-P7 represent partial blocks, where only the natural state (SE:SOF:SS/TS, TE:TOF:SS/TS) was used (Figure 4). The right side of the table describes which blocks addressed which predictions and which blocks were unusable due to a lack of viable embryos from one female.

| # of eyed egg embryos available |  |  |  |  |  |  | What predictions could block answer |  |  |  |
| --- | --- | --- | --- | --- | --- | --- | --- | --- | --- | --- |
| Block | SE:SOF:SS/TS | SE:TOF:SS/TS | SE:W:SS/TS | TE:W:SS/TS | TE:SOF:SS/TS | TE:TOF:SS/TS | Prediction 1A | Prediction 1B | Prediction 2A | Prediction 2B |
| **1** | 99 | 65 | 92 | 103 | 102 | 107 | **1A, 1B, 2A (no sperm swimming data)** |  |  |  |
| **2** | 133 | 129 | 119 | 131 | 141 | 140 |  |  |  |  |
| **3** | 134 | 130 | 130 | 2 | 2 | 3 | **Unusable, insufficient trout embryos** |  |  |  |
| **4** | 65 | 47 | 64 | 98 | 104 | 87 | **1A, 1B, 2A, 2B** |  |  |  |
| **5** | 131 | 137 | 94 | 21 | 26 | 32 |  |  |  |  |
| **6** | 96 | 94 | 91 | 136 | 142 | 144 |  |  |  |  |
| **7** | 51 | 72 | 37 | 109 | 106 | 124 |  |  |  |  |
| **8** | 141 | 132 | 116 | 96 | 103 | 109 |  |  |  |  |
| **9** | 95 | 97 | 103 | 136 | 144 | 143 |  |  |  |  |
| **10** | 101 | 99 | 104 | 138 | 125 | 148 |  |  |  |  |
| **P1** | 0 | N/A | N/A | N/A | N/A | 137 | **Unusable, insufficient salmon embryos** |  |  |  |
| **P2** | 123 | N/A | N/A | N/A | N/A | 48 | **1A, 1B** |  |  |  |
| **P3** | 55 | N/A | N/A | N/A | N/A | 89 |  |  |  |  |
| **P4** | 237 | N/A | N/A | N/A | N/A | 190 |  |  |  |  |
| **P5** | 276 | N/A | N/A | N/A | N/A | 103 |  |  |  |  |
| **P6** | 212 | N/A | N/A | N/A | N/A | 243 |  |  |  |  |
| **P7** | 101 | N/A | N/A | N/A | N/A | 124 |  |  |  |  |

**Appendix 2:**
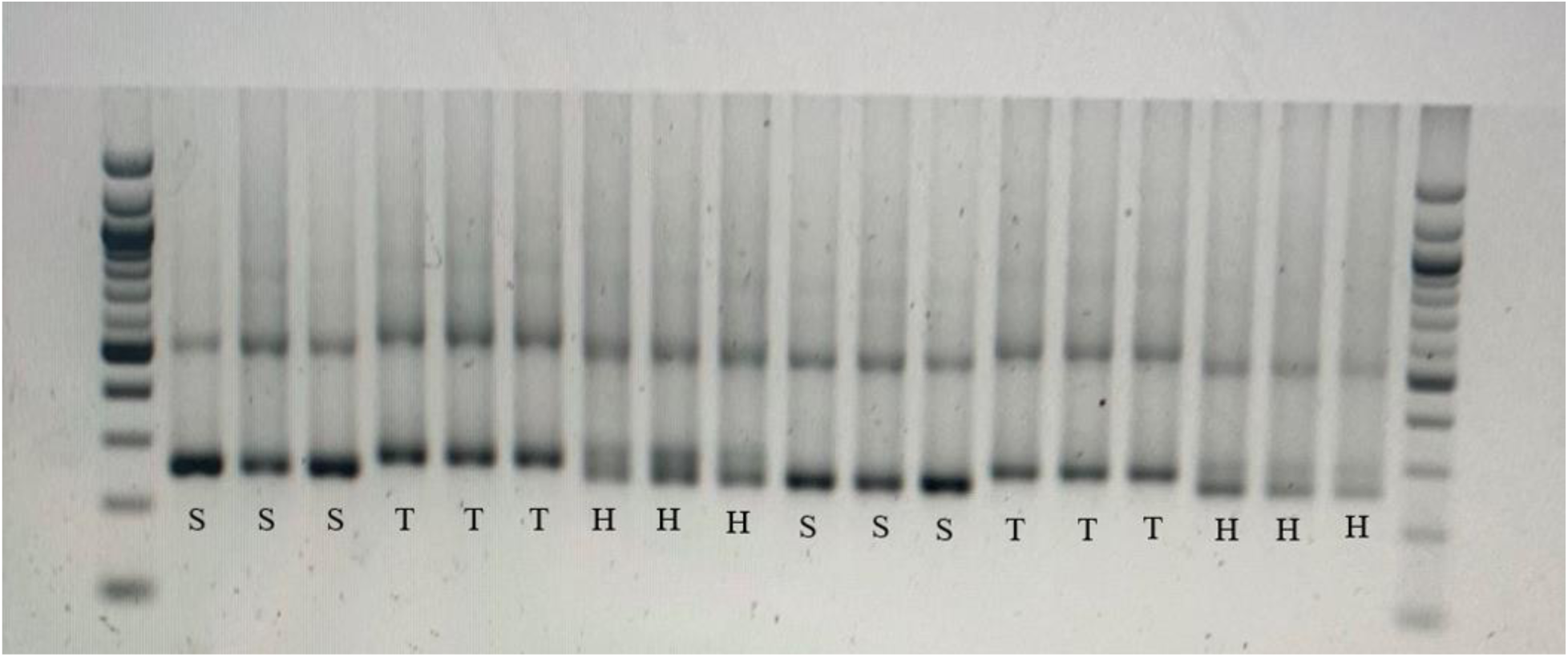
Agarose gel (2%) after electrophoresis using DNA from known Atlantic salmon (S), brown trout (T), and hybrids (H) from both salmon and trout mothers. Fish were taken from isolated wild populations, and hybrids produced from isolated in vitro fertilizations (without sperm competition). Pure salmon and trout display one bar, with trout bars being slightly higher. Hybrids display two bars, representing both species. DNA ladders are shown for comparison on each end of the gel.

**Appendix 3:** Number of embryos genotyped and used in paternity ratios per treatment in 15 blocks (1878 embryos total). Blocks 1-10 were full blocks and contain all treatments, but treatments TE:W:TS/SS and SE:W:TS/SS were not genotyped in the first two blocks due to not having sperm swimming data. Blocks P2-P7 are partial blocks and only tested the natural state (TE:TOF:TS/SS, SE:SOF:TS/SS). Embryos were genotyped in batches of six from each group at a time. The exact number used was based on whether paternity was extremely one-sided. A minimum of 18 embryos was used if one species consistently obtained a large majority (≥88%) of embryos within the first three batches, with 48 being the maximum number of embryos genotyped. Some groups had more than 18 embryos genotyped even when a majority was achieved.

| # of embryos genotyped and identified to father species (Salmon, Trout) |  |  |  |  |  |  |
| --- | --- | --- | --- | --- | --- | --- |
| Block | SE:SOF:SS/TS | SE:TOF:SS/TS | SE:W:SS/TS | TE:W:SS/TS | TE:SOF:SS/TS | TE:TOF:SS/TS |
| 1 | 24(24,0) | 48(29,19) | N/A | N/A | 36(28,8) | 48(26,22) |
| 2 | 48(12,36) | 30(6,24) | N/A | N/A | 30(7,23) | 24(0,24) |
| 4 | 24(24,0) | 36(8,28) | 48(20,28) | 18(1,17) | 48(32,16) | 48(27,21) |
| 5 | 24(0,24) | 24(3,21) | 18(0,18) | 18(0,18) | 18(16,2) | 24(0,24) |
| 6 | 48(30,18) | 30(8,22) | 18(0,18) | 18(0,18) | 30(22,8) | 48(13,35) |
| 7 | 24(24,0) | 18(18,0) | 18(16,2) | 24(20,4) | 30(22,8) | 30(24,6) |
| 8 | 30(7,23) | 18(2,16) | 48(14,34) | 18(1,17) | 24(3,21) | 48(14,34) |
| 9 | 48(24,24) | 48(32,16) | 18(2,16) | 18(2,16) | 18(2,16) | 24(1,23) |
| 10 | 48(11,37) | 30(6,24) | 18(0,18) | 18(1,17) | 18(0,18) | 24(0,24) |
| P2 | 48(35,13) | N/A | N/A | N/A | N/A | 18(16,2) |
| P3 | 18(16,2) | N/A | N/A | N/A | N/A | 48(38,10) |
| P4 | 30(26,4) | N/A | N/A | N/A | N/A | 18(0,18) |
| P5 | 48(16,32) | N/A | N/A | N/A | N/A | 18(0,18) |
| P6 | 48(13,35) | N/A | N/A | N/A | N/A | 30(6,24) |
| P7 | 18(17,1) | N/A | N/A | N/A | N/A | 48(14,34) |

**Appendix 4:**
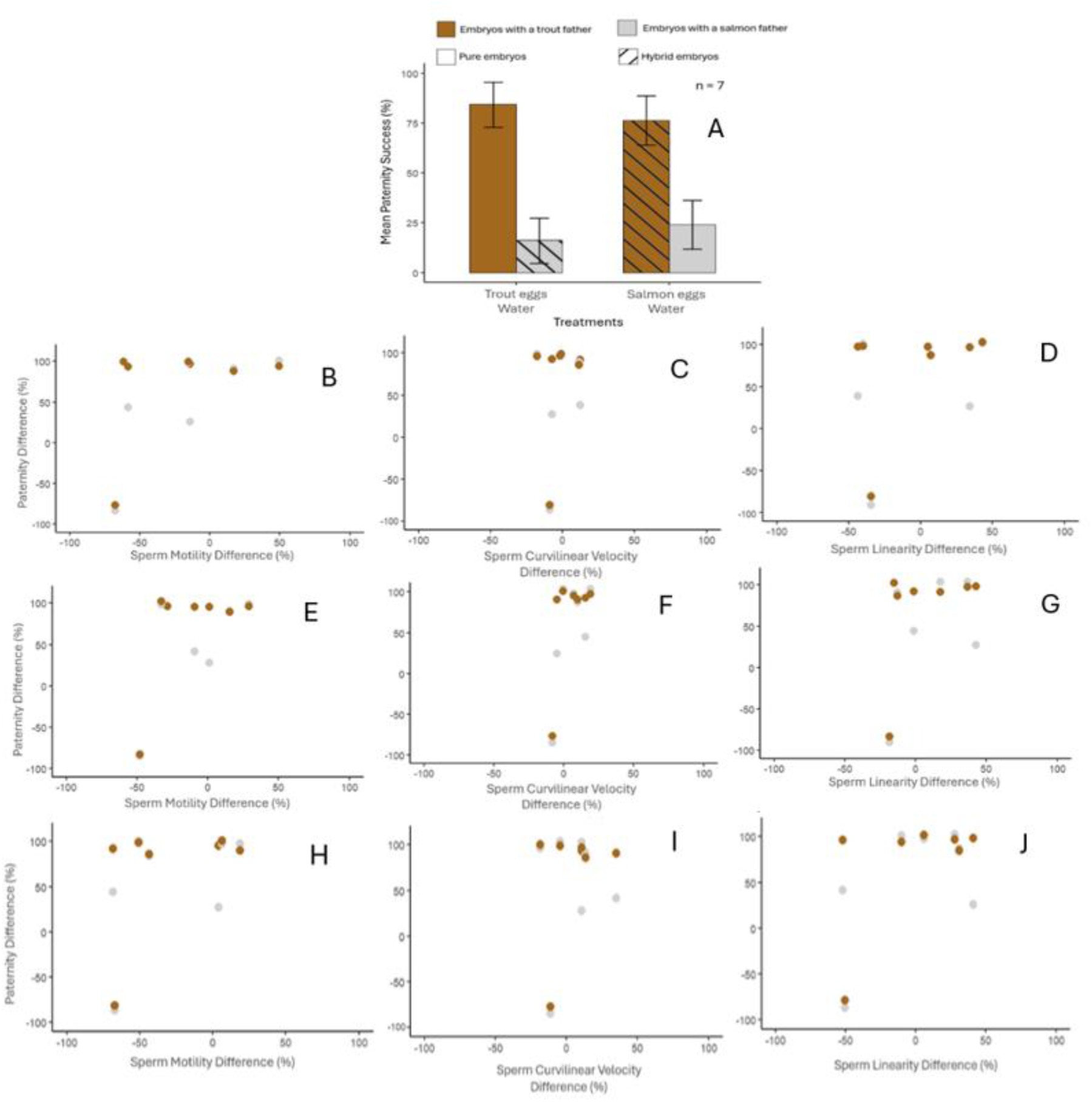
Mean paternity success across blocks between trout-salmon males under sperm competition without the presence of ovarian fluid (TE:W:SS/TS, SE:W:SS/TS) in both species of female. Hybrid embryos are represented by hatched bars. Relative paternity between males in each group sums to 100. Error bars represent standard error among 7 blocks (28 adult fish). Panels B, E, and H (percent of sperm cells that were motile), C, F, and I (sperm swimming velocity) and D, G, and J (sperm swimming linearity) compare the difference in paternity to the difference in semen quality between trout-salmon males within each block (e.g. if trout paternity is 30% lower than salmon paternity the paternity difference is −30). Semen quality measurements for each male were taken from performance in salmon ovarian fluid (panels B, C, and D), trout ovarian fluid (E, F, and G) and no ovarian fluid (H, I, and J) We fit linear mixed effects models for our treatments that did not contain ovarian fluid (salmon eggs in water; trout eggs in water) in the seven blocks (14 males; 14 females) that had sperm swimming performance data. The response variable was the difference in paternity between males, with either the difference in MOT, VCL, or LIN between males, as well as egg species and their interaction as fixed factors and block as a random factor (see equations 5-7) in each of our swimming mediums separately (SOF, TOF, W). No interactions between the difference in sperm quality and the species of the egg were found to be significant: In salmon ovarian fluid (MOT: ?² = 0.4497, df = 1, p = 0.5025, VCL: ?² = 0.1461, df = 1, p = 0.7023, LIN: ?² = 0.0029, df = 1, p = 0.9571) In trout ovarian fluid (MOT: ?² = 0.0206, df = 1, p = 0.8858, VCL: ?² = 0.1172, df = 1, p = 0.7321, LIN: ?² = 0.8057, df = 1, p = 0.3694) In water (MOT: ?² = 0.0127, df = 1, p = 0.9102, VCL: ?² = 2.4121, df = 1, p = 0.1204, LIN: ?² = 0.0078, df = 1, p = 0.9297). Within a block relative semen quality of each male did not clearly explain relative paternity: In salmon ovarian fluid (MOT: ?² = 1.9655, df = 1, p = 0.1609, VCL: ?² = 0.1476, df = 1, p = 0.7009, LIN: ?² = 0.7131, df = 1, p = 0.3984) In trout ovarian fluid (MOT: ?² = 1.8611, df = 1, p = 0.1725, VCL: ?² = 3.0550, df = 1, p = 0.0805, LIN: ?² = 0.4804, df = 1, p = 0.4883) In water (MOT: ?² = 1.2052, df = 1, p = 0.2723, VCL: ?² = 0.0698, df = 1, p = 0.7917, LIN: ?² = 2.4138, df = 1, p = 0.1203) Residuals were checked using Q-Q plots and normality was observed. All linear mixed effects models were fitted using the “lme4” package (Bates, 2007).

